# STRategy: A support system for collecting and analyzing short tandem repeats for forensic science

**DOI:** 10.1101/2023.02.20.529208

**Authors:** Nuttachai Kulthammanit, Poonyapat Sukawutthiya, Hasnee Noh, Kornkiat Vongpaisarnsin, Duangdao Wichadakul

## Abstract

Short tandem repeats (STRs) are short repeated sequences commonly found in the human genome. They provide many advantages to forensic sciences, such as identifying individuals, estimating the likelihood of kinship, and analyzing mixtures. Next-generation sequencing (NGS) technologies, e.g., ForenSeq Signature Prep, have been proposed for sequencing STRs, obtaining the sequence of each locus and SNPs, and inferring length-based alleles. However, even though the sequenced STRs from ForenSeq offer more insights into the STRs, which lead to the genetic analysis of population and sub-population structures, no open-source software platform enables the collection and management of STR data from NGS and incorporates related analysis tools in one place.

Here, we introduce STRategy, a standalone web-based application supporting essential STR data management and analysis capabilities. The analyzed data will be visualized in various forms, for example, charts, maps, and pattern alignments. The system implemented a role-based access control that allows users to search or access specific data depending on their responsibilities. It enables public users to search for data. In addition, they can view statistical data, for example, detailed alleles and genetic variation. Lab users can add, update, and see the information of individuals and explore pattern alignments for a specific locus within the population. Administrators can customize the system, for example, configure maps according to the samples’ geographic data, and manage reference STR repeat motifs.

We designed and developed the STRategy using software engineering principles for flexible extension and easy deployment utilizing the Docker container. The source code is publicly available at https://github.com/cucpbioinfo/STRategy. Also, we deployed a showcase system on a cloud computing service where its URL is included on the GitHub repository. The current version only supports the ForenSeq sample detail report files.

## Introduction

Short tandem repeats or short repeated Deoxyribonucleic acid (DNA) sequences are important DNA markers used in forensic fields. In addition, they have also been a biological marker for human individualization and relatedness. The STRs are advantaged in criminal cases [1, 2] due to the chance to figure out the DNA matches between questioned DNA profiles in DNA databases. Moreover, DNA databases are rapidly growing worldwide. Hence, the efficient management of STR data usable for criminal work investigation and population genetic studies is indispensable.

There are two popular techniques to investigate STR data: capillary electrophoresis (CE) [3] and next-generation sequencing (NGS) [4], which provide allele identification. However, NGS delivers more sequencing details than CE. Therefore, handling and analyzing STR data from NGS is less scalable and more challenging. To accelerate the use of sequence-based STRs from NGS for forensic science, in this paper, we propose the STRategy, a web-based application aiming to give convenience for collecting and analyzing the sequence-based STR data from ForenSeq sample detail report [5]. The system acts as an in-house database with a management system for STR data. Furthermore, it provides essential analyzing tools to analyze STR data in the database and visualizations to display results.

## Materials and Methods

### Overview

We designed the STRategy as a web-based application that allows users to organize and analyze short tandem repeat data efficiently and conveniently. The following subsections describe the underlying data and core methods designed and developed within the system.

### STR data

The STRategy accepts STR data generated from the ForenSeq Universal Analysis Software v1.3 called ForenSeq sample detail report; an Excel file with nine sheets, including Autosomal STRs, Autosomal STR Figure, Y STRs, Y STR Figure, X STRs, X STR Figure, iSNPs, iSNP Figure, and Settings described as follows.

- Autosomal STRs, Y STRs, and X STRs sheets - contain STR information for each chromosome, including tables of STR genotypes and allele sequence information for each locus (Figure 1). The STRategy will focus on the length-based allele table (Figure 1a) and sequence-based allele table (Figure 1b). Each locus will have a genotype in the length-based allele table derived from the sequences of that genotype in the sequence-based allele table with the value “Yes” in the “Typed Allele” column. For example, the locus TPOX (row 17 of Figure 1a) having genotype 8,11 in the length-based allele table has sequence detail from rows 54 and 57 of Figure 1b in the sequence-based allele table.
- iSNPs sheet - contains tables of single nucleotide polymorphisms (SNP) genotypes and SNP allele information for each locus (Supplementary Figure S1). This sheet resembles Autosomal STRs, Y STRs, and X STRs. The first table is the SNP genotype, and the second is the SNP allele information. The STRategy will focus on only these two tables.
- Autosomal STR Figure, Y STR Figure, X STR Figure, and iSNPs Figure - contain a summary table of read counts for each locus and a bar chart representing the overall read count. The STRategy does not collect data from these four sheets.
- Settings - contains all the constants used in the experiment. The STRategy also does not collect any information from this sheet.

**Figure 1.**
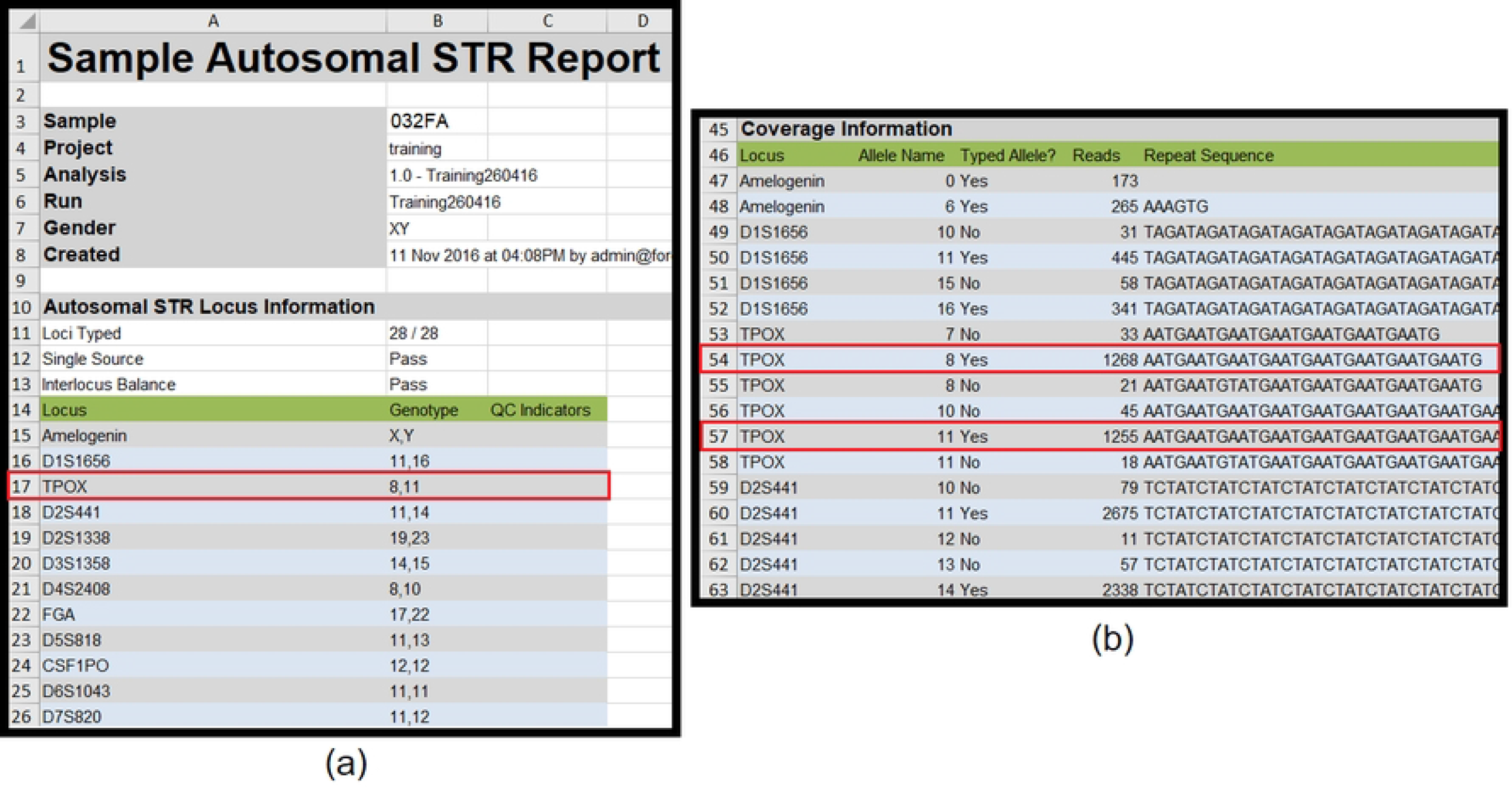
Autosomal STR data sheet from next-generation sequencing (ForenSeq) (a) STR genotype and (b) STR allele sequence information

STR data in ForenSeq sample detail report usually does not have a flanking region. There is another file called flanking report, which is not used by the system. However, some markers have flanking regions in the sample details report—for example, D1S1656 and D5S818. This file does not include personal information, for example, gender and race. All samples’ STR data are saved in the database with sample IDs that will be automatically linked to personal data when uploaded into the system by lab users.

The showcase STRategy contains 125 mocked samples constructed by swapping genotypes in the same locus among 125 actual ForenSeq sample detail reports. Note that the associated rows in the sequence-based allele table also moved with their swapped genotypes.

### Searching Module

The STRategy provides two sorts of searches, the Overview Search and Profile Search. The Overview Search utilizes the set of genotypes and loci to query and return the number of matched samples for public users and the sample IDs, countries, and provinces for lab users. The Profile Search requires a specific set of genotypes and loci and provides a correlation with statistical data. For the Overview Search, the system will search for samples that consist of a set of loci and genotypes specified by the user. For example, the user defines the loci and genotypes as D12S391:19,25 and TPOX:8,8. The algorithm then looks for samples that must include these loci and genotypes.

On the other hand, the Profile Search behaves similarly to OmniPop200.1 [6]. The user must first pick the core loci as the reference. After that, the user must specify the search input with minimal loci and genotypes of the sample according to the selected core loci. For example, if the user chooses core loci as European, the minimal loci are FGA, TH01, VWA, D1S1656, D2S441, D3S1358, D8S1179, D10S1248, D12S391, D18S51, D21S11, and D22S1045. With the input genotypes of the core loci, the STRategy will use Equations (1) and (2) to calculate genotype frequencies from allele frequencies, where Equations (1) and (2) are for homozygous and heterozygous genotypes, respectively. Then, the algorithm will multiply the reciprocal genotype frequencies from the same country and rank these values across the countries. The lowest value means the input profile is most common in that country.

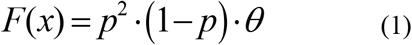

Where *p* is the allele frequency and

*θ* is a constant set by the lab user

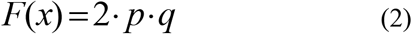

Where *p* is the first allele frequency and

*q* is the second allele frequency

### STR Pattern Alignment

The STRategy allows administrators to define the reference STR repeat motifs for each locus. This setting is in an Excel file, and an administrator has to upload it into the system. The file includes the locus name, the reference STR repeat motifs formatted according to the conventions used in [7], the allele, and the orientation columns (Supplementary Figure S2). The allele column with the “Default” value indicates that the pattern applies to all alleles in this locus. However, suppose the administrator wants the system to analyze an allele with a microvariant pattern. In that case, the administrator must add this microvariant explicitly, as shown in allele 10.3 of locus D2S441 (Supplementary Figure S2), which has a pattern different from other alleles within the same locus. Finally, the orientation column indicates the order of the samples’ sequence. The sequences in each locus of a sample details report file can be either forwarded or reversed. STRategy states that the reference STR repeat motifs must always be the forward strand (the sequence in the 5’ to 3’ direction). The administrator can set reverse or forward in the orientation column to handle the strand direction of samples’ sequences. The reverse orientation tells the system to reverse complement the reference STR repeat motifs before analyzing pattern alignment. For example, the reference STR repeat motifs of CSF1PO (row 3 in Supplementary Figure S2) is [ATCT]n in the forward strand. However, the administrator notices sequences in sample details report files are in the reverse strand direction. Hence, the administrator must set “Reverse” value in the orientation column. Then, based on these STR repeat unit patterns, the administrator can generate pattern alignment using a tool available on the administrator’s menu. This tool compares the reference STR repeat motifs of the locus to the DNA sequence within each sample locus and detects any insertion or deletion with a length less than the length of the reference motif.

### User management

The STRategy system provides three roles: public, lab, and administrator. Each role will have one of the four statuses: NOT_ACCEPTED, ACCEPTED, BLOCKED, and DELETED. For example, the status of a new user not yet approved by the administrator will be NOT_ACCEPTED. After approval, it will change to ACCEPTED. A user cannot access the system anymore if the administrator changes his/her status to BLOCKED or DELETED.

## Results

### Search Module

The STRategy provides three search options. The first option allows users to upload a sample file as input (Figure 2a), which is then searched against all available samples within the system. The second and third options are equivalent in using a set of genotypes and loci as the input. However, the second option (Figure 2b) allows an independent set of loci, while the third option (Figure 2c) allows only the loci corresponding to the selected kit. All options use the input genotypes and loci to search against all samples within the system and return only the number of matched samples for public users (Figure 2d). Lab users will also get additional information, including the number of matched samples grouped by geographical location, sample IDs, nations, and provinces (Figure 2e).

**Figure 2.**
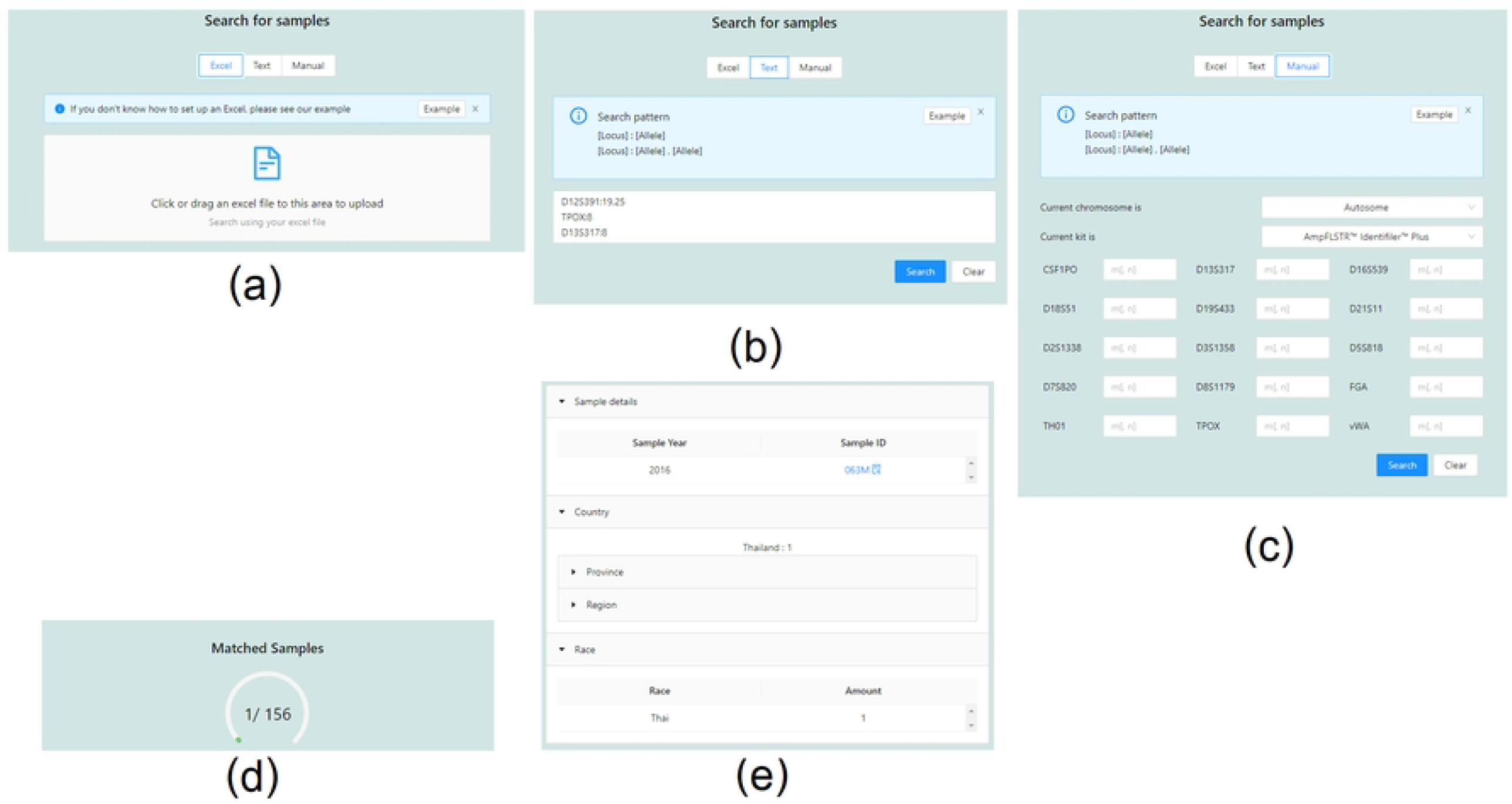
Overview Search options. (a) search by ForenSeq sample detail report files of a sample (b) search by any loci and genotypes (c) search by any loci and genotypes of a specific test kit (d) search results for public users (e) additional search results for lab users

The Profile Search (Figure 3a) lists the required loci called “Core STR Loci.” The system obtained the core STR loci used for human identification from STRBase [8] and length-based allele frequency data from STRidER [9, 10] as defaults for the correlation calculation. After selecting core loci, the user must provide genotypes for each required locus. There are two methods to input genotypes. The first method is to enter genotypes manually. The second method is to upload an input file, automatically entering each core locus’s genotype fields. The underlying algorithm will then use these input genotypes to calculate the correlation of the sample with each country. The system then displays the results with a table of the countries sorted by the most common genotype frequencies (Figure 3b) and detailed tables of the length-based allele frequencies calculated from each country (Figure 3c). The administrator can customize the core STR loci and the length-based allele frequency data (See section system management by administrators).

**Figure 3.**
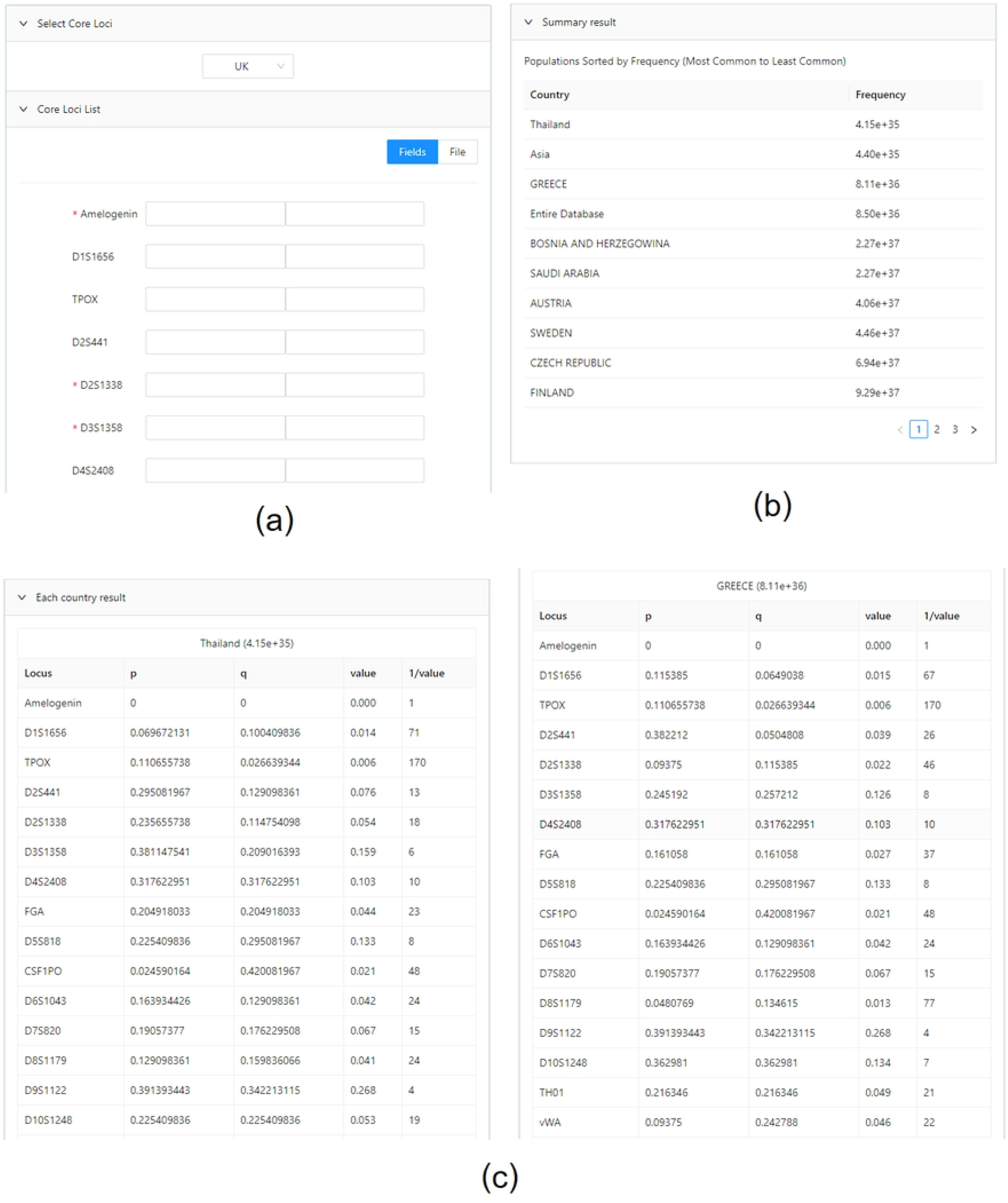
Profile Search. (a) input form (b) countries with most common genotype frequencies (c) detailed genotype frequencies of each country and locus

### Statistics

The system analyzes the STR data in the database in several ways. First, it shows the results in various forms so that users may interactively explore the results via the web pages. In addition, the system allows public users to access only aggregated data, while lab users can also access individual data. This section describes the statistical data analyzed by the system.

The Genetic variation (Figure 4) and Allele details (Figure 5) pages display the allele frequencies for each locus. Users may pick autosomal, X, or Y STRs and a locus on the left side of the web page (Figure 4a). Selecting a locus will show the bar graph of length-based allele frequencies of that locus, along with its statistical values on the right-hand side (Figure 4b), divided into three cases. First, the locus is composed of haploid genotypes only (Supplementary Figure S3). Therefore, the system will display the Match Probability (PM), Polymorphic Information Content (PIC) [11], and Power of Discrimination (PD) values. Second, the system will display the Observed Heterozygosity (Hobs), Expected Heterozygosity (Hexp), Match Probability (PM), Polymorphic Information Content (PIC), Power of Discrimination (PD), and Power of Exclusion (PE) if the locus contains diploid genotypes only (Supplementary Figure S4). Third, if the locus contains both haploid and diploid genotypes, for example, a locus of the X chromosome, the system will allow the user to toggle between the Diploid and Haploid (Figure 4c). Figure 4d displays a table containing all sequence-based allele variations when the user clicks an allele bar from the bar chart in Figure 4b. Additionally, the allele frequencies summarized from all samples are available on the Allele details page (Figure 5), divided into chromosomes, STR variations, and allele frequencies for each variation.

**Figure 4.**
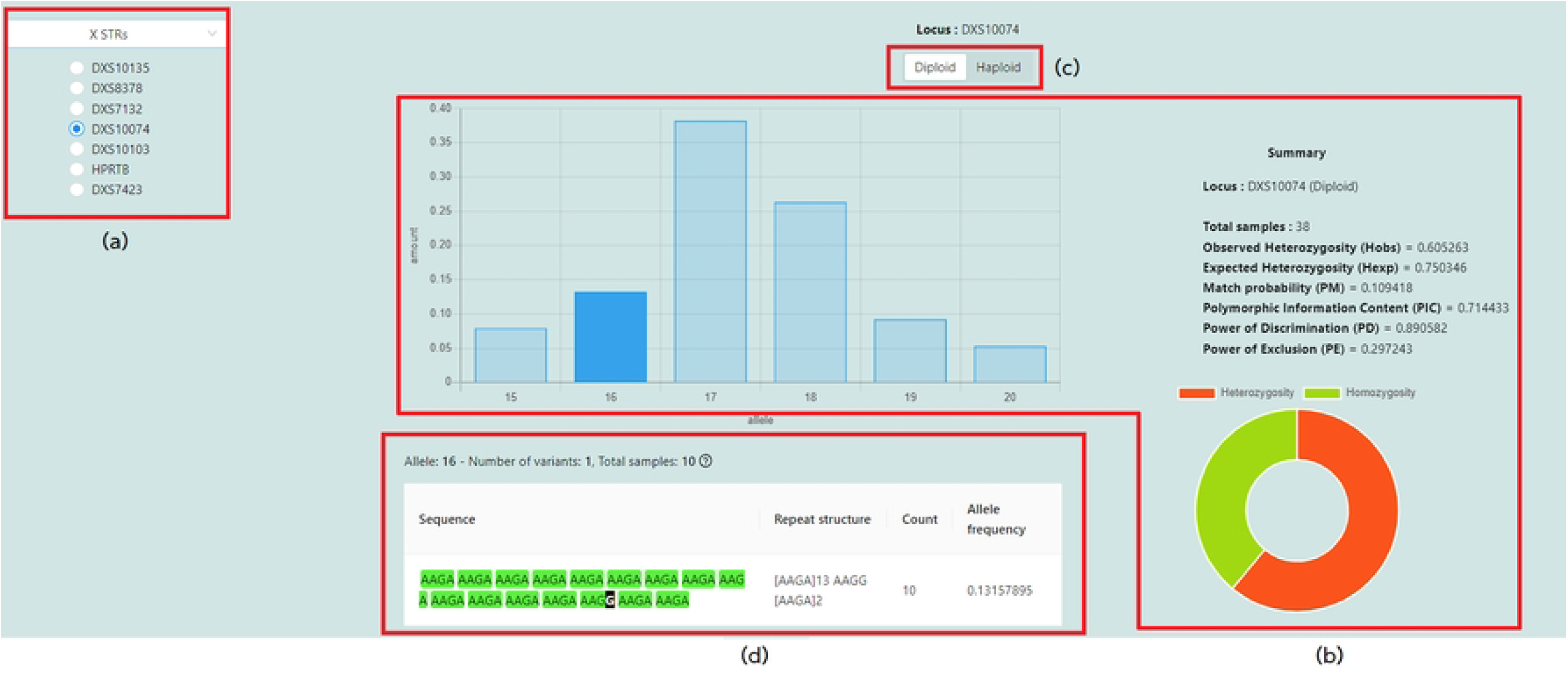
Genetic variation page. (a) list of selectable chromosome and loci (b) allele frequencies and common statistical values for assessing genetic variation of the samples (c) diploid or haploid toggle, and (d) genetic variation of the selected locus DXS10074 with allele 16

**Figure 5.**
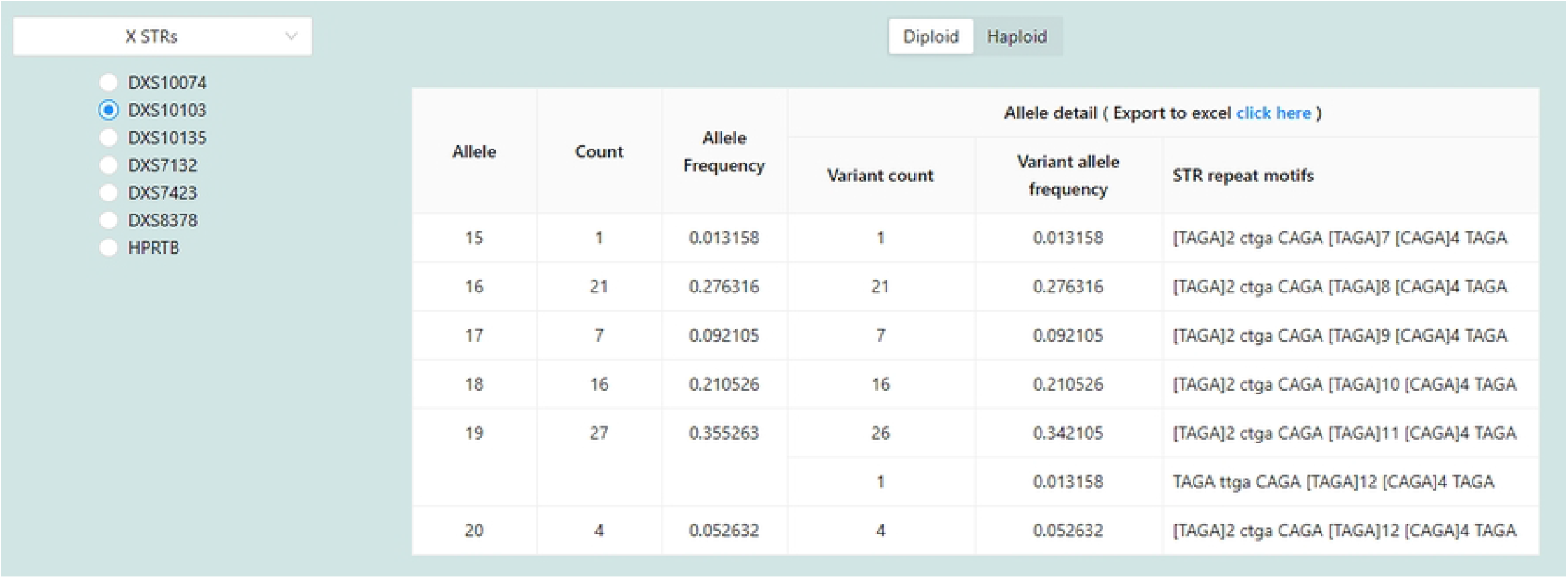
Allele details page.

Allele distribution by geographic page shows the map of samples with regional data. Each circle on the map represents the number of samples. Hovering the cursor over a circle will display all length-based alleles found within the province (Figure 6). Additionally, the user may mouse over a square to see the number of samples collected from each region. This map can be changed to a different country (See section system management by administrators).

**Figure 6.**
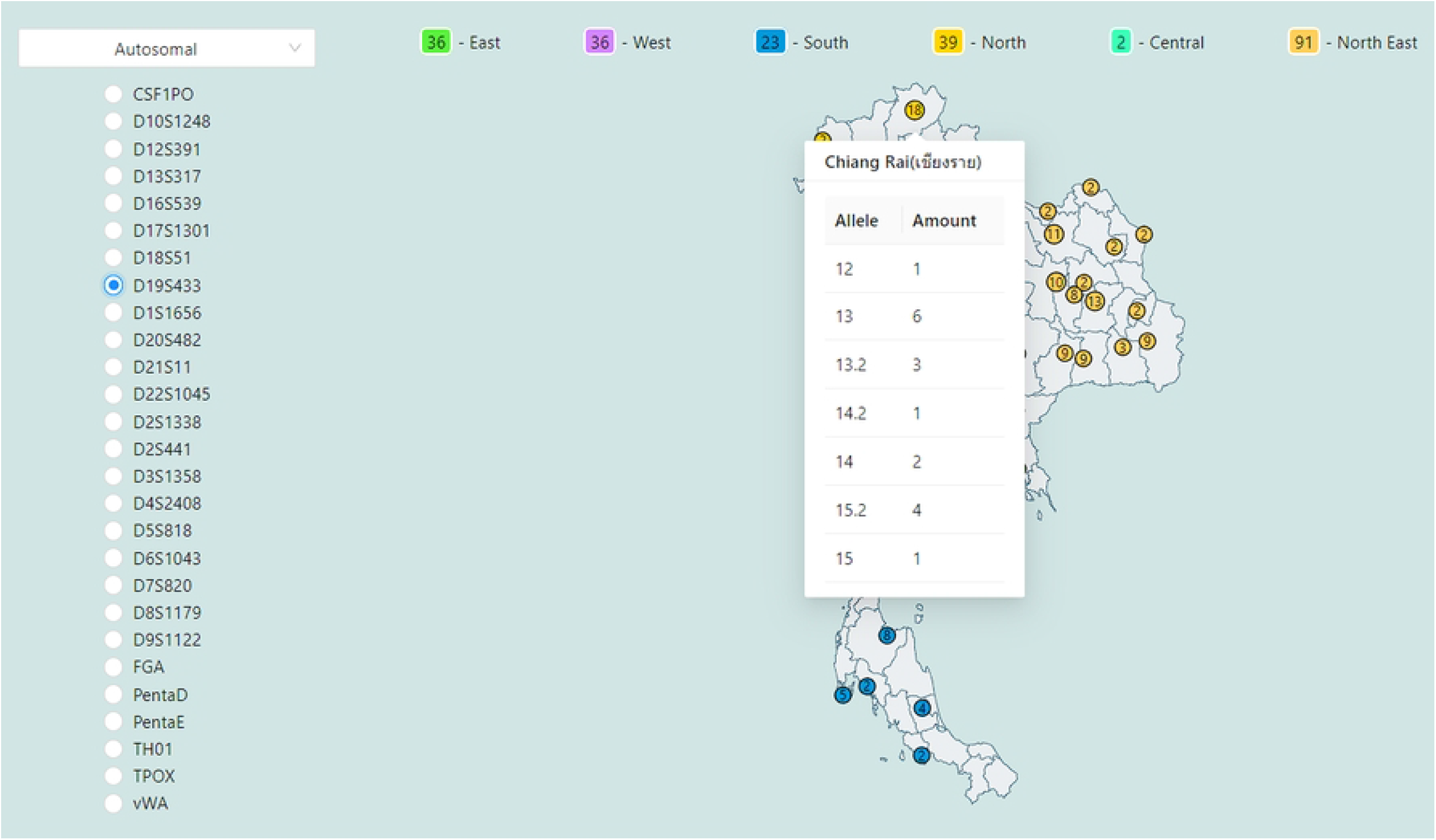
Allele distribution by geographic location of a selected locus.

The iSNP Statistic Summary (Supplementary Figure S5) page displays data aggregated by SNP locus and allows for searching by locus name. The bar graph illustrates the summary of the altered nucleotide pairs of all samples within the database. The mouse hovering over each bar shows the proportion of each SNP variation.

The allele frequency comparison page (Supplementary Figure S6) provides a comparative view of length-based allele frequency from all samples within STRategy’s local database and international populations obtained from STRidER [9]. Users can explore the allele frequency of a specific locus by selecting from the drop-down list. The system will then generate a two-dimensional grid in which each row represents the allele, and each column indicates the country. Each cell represents the allele frequency by color and shows the value when mouse over.

### STR pattern alignment

The STR pattern alignment page allows users to browse through chromosomes, loci, and alleles. After selection, it will display tabular data in five columns: Sample Year, Sample ID, Sequence, Read count, and STR repeat motifs, where the Sequence column contains the sample nucleotide sequence with each STR motif highlighted individually (

Figure **7**) and the STR repeat motifs column includes a summary of the sequence’s STR. This data is not accessible to public users. Only lab users and administrators can access this page.

Figure **7** shows the pattern alignment of all samples within the database on the locus FGA with allele 24.2. The system displays the reference STR repeat motifs on the top of the pattern alignment table, telling the user that its pattern within the sequence is reverse or forward as pre-specified by the administrator in Supplementary Figure 2. When the administrator uploads a sample details report file to the system where sequences in the CSF1PO locus are reversed, the administrator must specify the pattern of this locus as the reverse in the orientation column, and the STRategy will display sequences on this page as reverse. On the contrary, the sequences from the locus D10S1248 in the sample details report file are forwarded; hence, the orientation column of this locus must be pre-specified as forward. So, the system will display them as forward. However, users can view the forward or reverse directions by toggling the referenced pattern tag or kebab menu. For example, Figure 7 shows that the reference STR repeat motifs are reversed. It means the original sequences of this locus within the sample details report file are in the reverse direction. On the other hand, if users toggle the referenced pattern, all sequences will be forward, as shown in Supplementary Figure S7.

**Figure 7.**
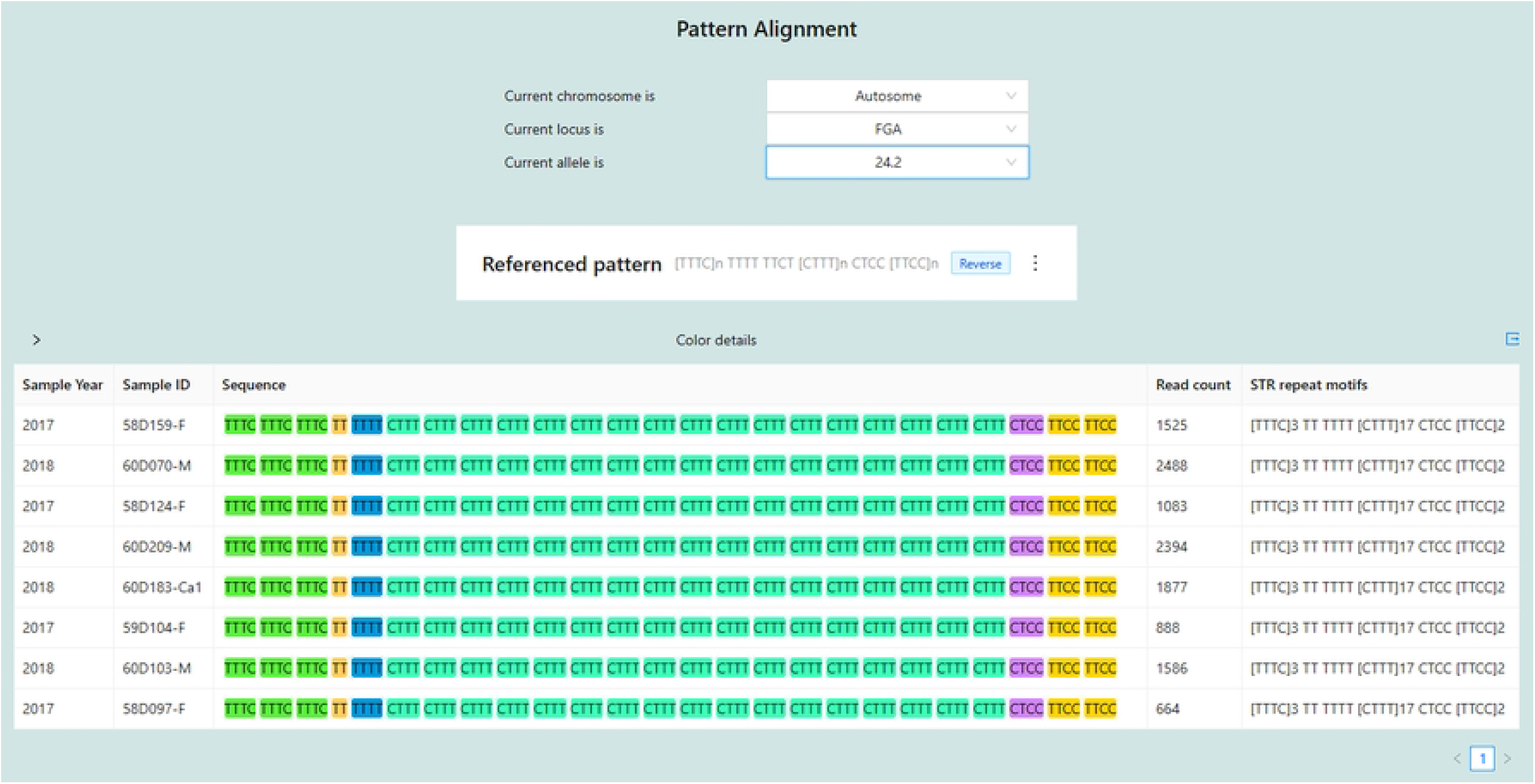
Pattern alignment page.

Lab users can also export the pattern alignment results in Excel formats, i.e., XLS and XLSX extensions, via the export page (Supplementary Figure S8). Supplementary Figure S9 shows an example result of the exported alignment patterns with five columns: Sample ID, Sample Year, Allele, Repeat Structure, and Sequence.

### Data management by lab users

Lab users can access all functions as public users. Lab users can also upload STR ForenSeq sample detail report files to the system via the upload STR data page (Supplementary Figure S10). The system assumes that the lab users have validated the files following the ForenSeq manual and their data validation protocol. When the lab user uploads the file, the system checks the sample ID to see whether it duplicates a sample ID already in the system. If this is the case, the system offers the user the choice of canceling or overwriting the sample data (Supplementary Figure S11). After uploading sample details report files, lab users can also upload the corresponding personal data file (Figure 8) for each sample or as a batch of persons via the upload personal data page (Supplementary Figure S12). The system links these personal data based on the sample ID. Lab users can edit or delete personal data via the manage personal data page (Supplementary Figure S13). The system extracts the value for each column as follows. The gender can only be MALE or FEMALE. The system will then look for values in the Province, Region, Country, and Race columns and associate them with the sample ID. An administrator must enter these provinces, regions, countries, and races into the system before a lab user uploads this personal data file.

**Figure 8.**
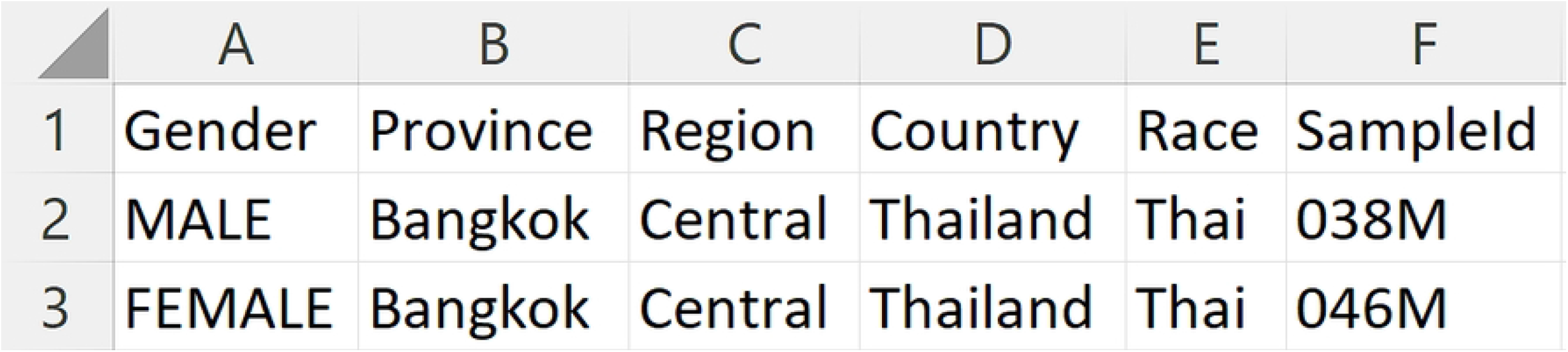
Personal information in Excel file format.

### System management by administrators

Administrators can perform the same functions as public users and lab users. Additionally, they can customize the system by, for example, changing the map and managing users (Supplementary Figure S14). The STRategy system offers the administrator’s default username and password for the initial login, whose password can be changed later.

Administrators may use any country’s map for map representation (Supplementary Figure S15) for visualizing allele distribution by geographic location (Figure 6). Moreover, they can modify the center of the map depending on the latitude and longitude grid system and the map amplification scale to accommodate the data in their local database. The map visualization was created using React Simple Maps [12] and supports data in TopoJson [13] or GeoJson [14] formats. However, the large map nation file will degrade the system’s performance. The display will be slower if the map file is large. There are examples in [15] of how to get and utilize the map in this system. If the developer wishes to generate customized TopoJson files based on shapefile data, a conversion guide is available in [16].

The system also provides web pages for setting the Profile Search. The first page is to upload the core loci to the system or replace the existing core loci with new core loci (Supplementary Figure S16). This file’s format is an Excel file consisting of two columns: locus and country (Supplementary Figure S17). The default core loci within the file will also appear as part of other countries’ core loci. For example, the default core loci are FGA, TH01, TPOX, and vWA, and a country ABC’s core loci are only FGA and TPOX. When displayed on the Profile Search, users will see all the default core loci: FGA, TH01, TPOX, and vWA, where FGA and TPOX will be marked as required. The second page is to upload length-based allele frequencies (Supplementary Figure S18). The STRategy has default length-based allele frequency data from STRidER [9]. However, the administrator can replace default data by uploading his/her Excel files. There are two upload options to replace length-based allele frequencies in the database. First, upload allele frequencies of all countries simultaneously with an Excel file, as shown in Supplementary Figure S19. This Excel file contains length-based allele frequency tables; each table represents the allele frequencies of a locus. All tables have a locus name on the top of the tables. The second option is uploading allele frequencies of all loci by country (Supplementary Figure S20). This Excel file does not have a country name, but the administrator must choose the country before uploading the file (Supplementary Figure S21). The last page (Supplementary Figure S22) is to set up the theta constant of Equation (2) (homozygous genotype equation).

Additionally, administrators can add new loci and kits (Supplementary Figure S23) to the system. They can also add and modify countries and create, update, and remove races (Supplementary Figure S24), regions, and provinces (Supplementary Figure S25) associated with each country (Supplementary Figure S26). Then, the geographic page uses this information to display the map correlated with the uploaded samples’ personal data (Figure 8). Finally, they may submit reference STR repeat motifs and regenerate repeat motifs of the samples for the Pattern alignment. When lab users submit new samples to the system or administrators upload new reference STR repeat motifs for pattern alignment, the system will inform administrators to conduct the regeneration for new pattern alignments (Supplementary Figure S27).

### Implementation

The STRategy frontend was implemented based on React library [17] version 16.3. React renders the virtual DOM, which works fast as it only changes individual DOM elements instead of reloading the complete DOM every time. Also, it provides reusable components making it easier for the developer to develop the system. We implemented the STRategy backend using Spring Boot [18], an open-source micro-framework built on top of the Spring framework, as the forensic data is sensitive to data privacy. The Spring framework provides industry-standard security schemes and delivers a trustworthy solution. The system uses MySQL database version 5.7 [19] to store all the data.

### Validation

We calculated the allele frequencies and statistical values using 125 samples imported into the STRategy as the showcase and compared them to the results from STRAF [20]. All five comparison results of allele frequencies and statistical values: autosomal (diploid), x (diploid), x (haploid), y (diploid), and y (haploid), arranged as five folders were compressed into the Supplementary File S1 (See Additional data section). Each folder has three sub-folders inside. The first folder contains allele frequency results. This folder has three files: results from STRAF (Supplementary Figure S28), results from STRategy (Supplementary Figure S29) and compared results from both (Supplementary Figure S30). The second folder contains statistical results with three files: results from STRAF (Supplementary Figure S31), results from STRategy (Supplementary Figure S32), and compared results from both (Supplementary Figure S33). The last folder contains raw data files uploaded to STRAF. We validated allele frequencies and statistical values to 5 decimal places for all loci listed in

Supplementary Table **S1**. For diploid, we compared STRategy and STRAF by five values: Polymorphic Information Content (PIC), Match Probability (PM), Power of Discrimination (PD), Heterozygosity (Hobs), and Power of Exclusion (PE). STRategy and STRAF, respectively, reference the formula from [22] and [23] to calculate gene diversity (GD). So, we did not compare GD results in this validation. For haploid, we measured STRAF and STRategy by three values: Polymorphic Information Content (PIC), Match Probability (PM), and Power of Discrimination (PD). The validation results show that all values calculated by STRategy are consistent with all values computed by STRAF.

We validated the Pattern alignment results from STRategy by comparing them with the results from STRait Razor 3.0 [21]. We randomly selected five samples’ pattern alignment from 125 samples generated by STRategy. After that, we used STRait Razor to get pattern alignment from FASTQ files of those five samples and imported the results files from STRait Razor into the Excel-based workbook of STRait Razor 3.0 [22] from Strait Razor’s GitHub [23]. This workbook will display pattern alignments for each sample. After receiving the STRategy and Strait Razor results, we compared every locus reported by STRait Razor (

Supplementary Table **S2**). The comparison results were saved to Excel files and compressed into Supplementary File S2 (See Additional data section). Moreover, our 125 examples have ten loci with indel alleles (

Supplementary Table **S3**). So, we randomly selected ten samples with those loci and compared the alignment results with Strait Razor. As STRait Razor does not have pattern alignment of some alleles in some loci, for example, allele 15.3 of locus D8S1179, we omitted the comparison of those alleles. Nevertheless, the validation results show that all pattern alignments generated by STRategy agreed with all pattern alignments identified by the Excel-based workbook of STRait Razor 3.0. Note that we focused on validating the repeat regions and not including flanking regions.

## Conclusion

The STRategy is a standalone web-based application facilitating various analyses and administration of short tandem repeat data. The system delivers essential data management capabilities and several visualizations to interactively explore the statistically summarized data. In addition, the STRategy provides an STR pattern alignment tool, which includes capabilities for identifying STR repeat motifs, locating STR variations, and assisting lab users in discovering new variants with allele details. Hence, the STRategy is a helpful system for managing and analyzing length-based and sequence-based STRs from NGS for forensic science. Finally, the STRategy provides features for an administrator to customize and extend the system flexibly, such as changing the map to different countries or adjusting the pattern alignment settings. Administrators can customize the STRategy for their laboratory. We plan to include STR files from CE and other NGS platforms in the next version.

## Acknowledgements

We want to thank the department of computer engineering, faculty of engineering, Chulalongkorn university for providing computing facilities.

## Supporting information

**Supplementary Figure S1 SNP data sheet from next-generation sequencing (ForenSeq)** (a) SNP genotype and (b) SNP allele information

**Supplementary Figure S2 Example of reference STR repeat-unit patterns in Excel file format**

**Supplementary Figure S3 Genetic variation page with only haploid genotypes under the statistics section**

**Supplementary Figure S4 Genetic variation page with only diploid genotypes**

**Supplementary Figure S5 iSNP statistical summary page under the statistics section**

**Supplementary Figure S6 STRategy and STRidER allele frequency comparison page of locus D3S1358 under the statistics section** (a) allele frequency calculated from all samples within the STRategy database (b) allele frequency of various countries obtained from STRidER

**Supplementary Figure S7 Pattern alignment page with forward reference pattern under the lab user section**

**Supplementary Figure S8 Pattern alignment export page under the lab user section**

**Supplementary Figure S9 An exported Excel file of the pattern alignment result**

**Supplementary Figure S10 Upload STR data page under the lab user section**

**Supplementary Figure S11 Lab user uploads duplicated data to the system**

**Supplementary Figure S12 Upload personal data page under the lab user section**

**Supplementary Figure S13 Manage personal data page under the lab user section**

**Supplementary Figure S14 Manage user page under the administrator section**

**Supplementary Figure S15 Manage map admin page under the administrator section** (a) Preview page (b) Configuration panel

**Supplementary Figure S16 Web page for managing core loci under the administrator section**

**Supplementary Figure S17 An example of Excel file for core loci upload**

**Supplementary Figure S18 Web page for uploading allele frequency of all countries under the administrator section**

**Supplementary Figure S19 Length-based allele frequencies of all countries**

**Supplementary Figure S20 Length-based allele frequencies of a country**

**Supplementary Figure S21 Web page for uploading allele frequency by a country**

**Supplementary Figure S22 Web page for managing theta constant under the administrator section**

**Supplementary Figure S23 Web page for managing kits and loci under the administrator section**

**Supplementary Figure S24 Web page for managing races under the administrator section**

**Supplementary Figure S25 Web page for managing provinces under the administrator section**

**Supplementary Figure S26 Web page for managing countries under the administrator section**

**Supplementary Figure S27 Notification to administrators for the management of pattern alignment under the administrator section**

**Supplementary Figure S28 Allele frequencies result file from STRAF (X diploid)**

**Supplementary Figure S29 Allele frequencies result file from STRategy (X diploid)**

**Supplementary Figure S30 Allele frequencies comparison result file from both STRAF and STRategy (X diploid)**

**Supplementary Figure S31 Statistical values result file from STRAF (X diploid)**

**Supplementary Figure S32 Statistical values result file from STRategy (X diploid)**

**Supplementary Figure S33 Statistical values comparison result file from both STRAF and STRategy (X diploid)**

**Supplementary File S1 Length-based allele verification**

**Supplementary File S2 Pattern alignment verification**

**Supplementary Table S1 loci list in allele frequencies and statistical values verification**

**Supplementary Table S2 loci list in pattern alignment verification**

**Supplementary Table S3 Additional pattern alignment loci**

